# Odor concentration change detectors in the Olfactory Bulb

**DOI:** 10.1101/114520

**Authors:** Ana Parabucki, Alexander Bizer, Genela Morris, Matthew C. Smear, Roman Shusterman

## Abstract

Dynamical changes in the environment strongly impact our perception ^1,2^. Consistent with this, sensory systems preferentially represent stimulus changes, enhancing temporal contrast ^3,4^. In olfaction, odor concentration changes across consecutive inhalations (ΔC_t_) can guide odor source localization. Yet the neural representation of ΔC_t_ has not been studied in vertebrates. We have found that a subset of mitral/tufted (M/T) cells in the olfactory bulb explicitly represent ΔC_t_. These concentration change detectors are direction selective: some respond to positive ΔC_t_, while others represent negative ΔC_t_. This change detection enhances the contrast between different concentrations and the magnitude of contrast enhancement scales with the size of the concentration step. Further, ΔC_t_ can be read out from the total spike count per sniff, unlike odor identity and intensity, which are represented by fast temporal spike patterns. Our results demonstrate that a subset of M/T cells explicitly represents ΔC_t_, providing a signal that may instruct navigational decisions in downstream olfactory circuits.

## Introduction

The brain must track how external information changes with time. Correspondingly, sensory circuits deploy specialized cell types for dynamic stimuli: visual neurons emphasize luminance changes and motion ^4^, auditory neurons capture amplitude and frequency modulation ^5^, and somatosensory neurons encode vibrating touches ^6^. Odor stimuli also change dynamically in ways that are relevant to odor source localization. Although odor source localization depends partly on comparison of odor concentration across the two nares, animals can still find odor sources and follow odor trails with one naris blocked ^7-9^. This remaining ability shows that animals also perform temporal comparison of odor concentration, from sniff to sniff (*ΔC_t_*), to guide them to an odor source. Yet despite this evidence that ΔC_t_ can guide odor tracking, whether olfactory neurons encode sniff to sniff changes has not been studied in vertebrates.

Unlike invertebrate olfactory systems ^10-15^, in which olfactory sensory neurons (OSNs) are continuously exposed to the medium, air-breathing vertebrates discretize the input to OSNs into intermittent inhalations. In this case, the brain must maintain a memory of odor concentration across the exhalation interval to compute ΔC_t_. How and where does the olfactory system solve this task? We demonstrate here that a subset of neurons in the olfactory bulb explicitly encode ΔC_t_ on the time scale of a single sniff. Thus, like their counterparts in other sensory systems, a subset of olfactory neurons specializes in representing stimulus dynamics.

## Results

### Experimental setup and response types

We recorded respiration and M/T cell activity (7 mice, 92 cells, 242 cell-odor pairs) in awake, head-fixed mice (Fig. 1a). To rapidly change odor concentration, we passed odorized airflow through a concentration change manifold (Fig. 1a, Methods). Sniffing was measured through an intranasal pressure cannula (Fig. 1a). Using real-time closed-loop odor presentation, we switched odor concentrations at the beginning of the exhalation phase so that the stimulus reached its new steady state concentration before the onset of the next inhalation (Fig. 1b, S1a).

**Figure 1.**
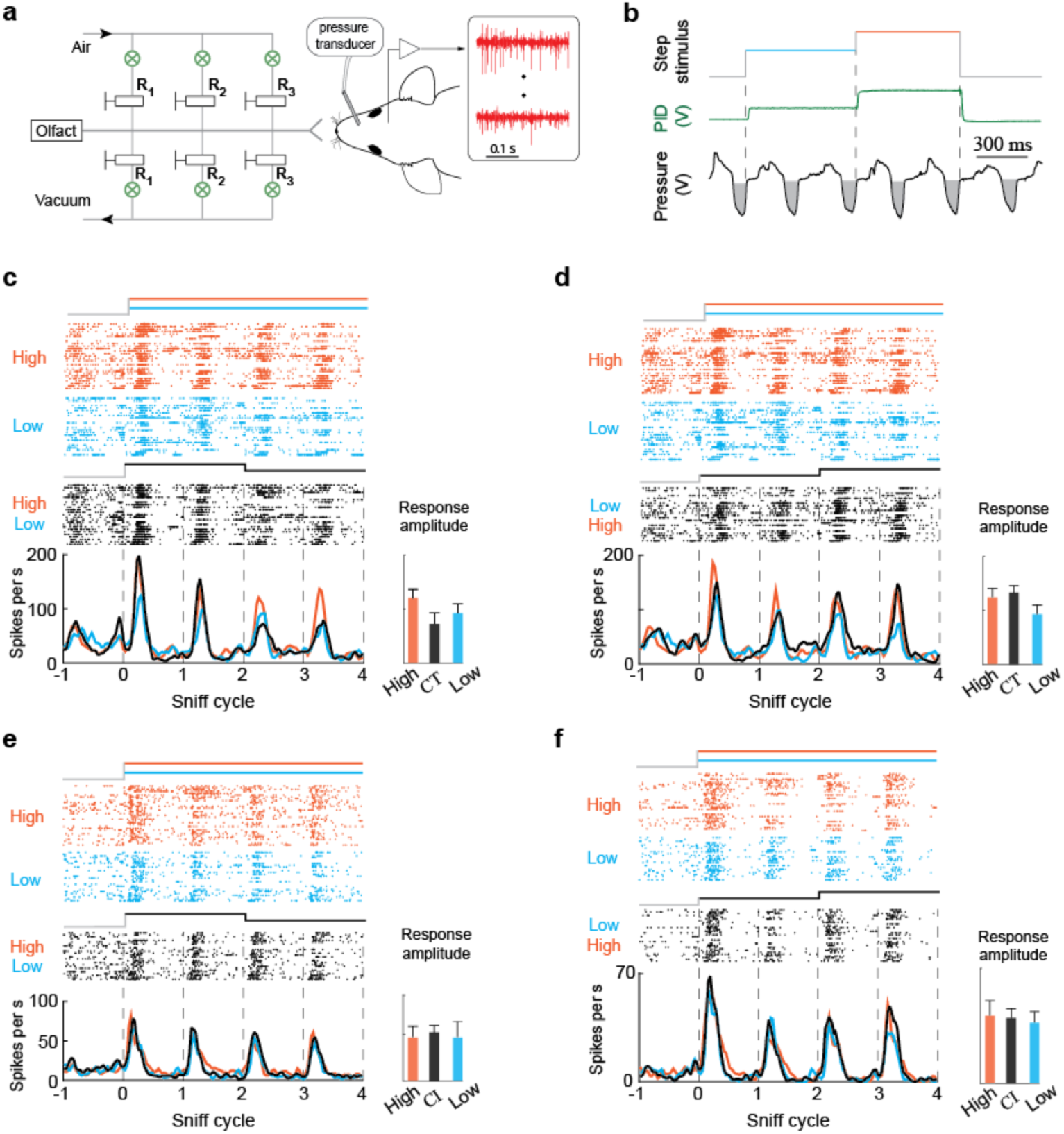
Concentration tracking and concentration invariant M/T cells. **a.** Schematic of the experiment. Right: A head-fixed mouse implanted with an intranasal cannula and a multielectrode chamber was positioned in front of the odor delivery port. Left: concentration change manifold. **b.** Odor concentration step paradigm. Odor concentration changes every two sniff cycles. Green curve indicates the response of a photoionization detector (PID) to presentation of ethyl acetate. Sniff waveforms (black) are shown below the plots. Grey areas indicate inhalation. Vertical dashed lines indicate onset of concentration changes. **c.-d.** Examples of concentration tracking responses. Raster and PSTH plots of M/T cell response to static high concentration (orange), static low concentration (blue) and concentration step stimuli (black). The responses of these cell odor pairs change with odor concentrations the same way in both static and step stimuli. Bar graph on right shows peak response amplitudes on the third sniff cycle for each stimulus. Error bars indicate standard deviation. **e.-f.** Same as **c-d**, but for cell-odor pairs that are invariant to odor concentration.

In the first set of experiments, we presented odorants in two static concentration patterns: high (H), low (L), and two dynamic patterns: a step from high to low (HL), and a step from low to high (LH). The high concentration was twice that of the low concentration. Step stimuli consisted of a presentation of one concentration for two sniff cycles, followed by a switch to the other concentration. These stimuli evoked three different response types across odor-cell pairs. Concentration-tracking (CT; Fig. 1c-d) responses exhibited different activity to high and low concentrations. The responses to the step stimuli were indistinguishable from responses to the static stimuli. These cell-odor pairs thus faithfully represent the concentration on each sniff. Another type of response is concentration-invariant (CI; Fig. 1e-f). These cell-odor pairs responded identically to both static stimuli, as well as to ΔC_t_ stimuli. These unchanging responses may be specialized for odor identification, for which concentration invariance is an important property ^16,17^. Alternatively, these cells may be in the saturated range of their concentration response function for this odor.

**Figure 2.**
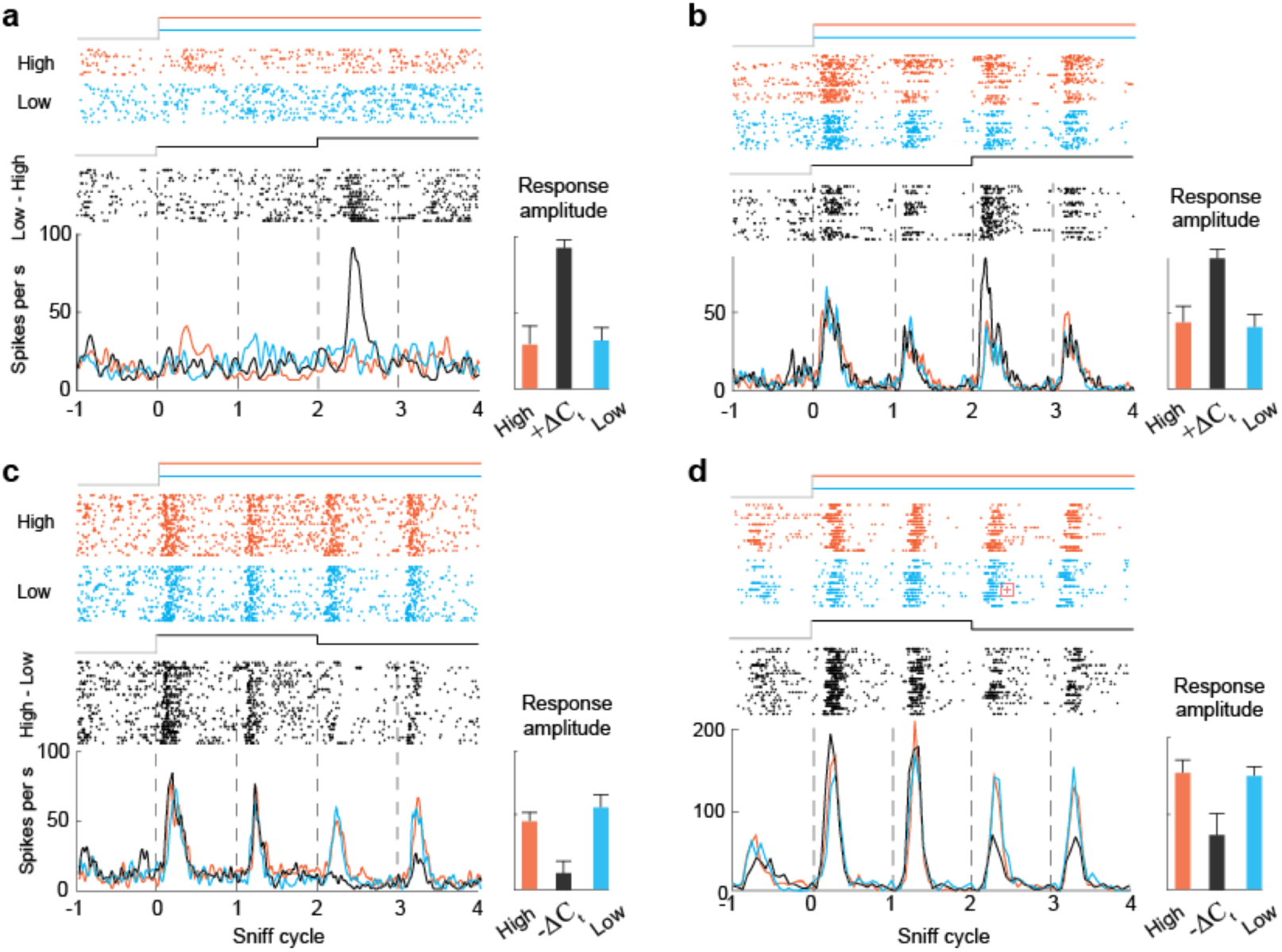
M/T cells responsive to changes in odor concentration. **a. b.** Examples of +ΔC_t_ responses. Raster and PSTH plots of M/T cell response to static high concentration (orange), static low concentration (blue) and low to high (black). Bar graph on right shows peak response amplitudes on the third sniff cycle for each stimulus. Error bars indicate standard deviation. **c,d.** Examples of -ΔC_t_ responses. Raster and PSTH plots of M/T cell response to static high concentration (orange), static low concentration (blue) and high to low stimulus (black).

Lastly, we observed responses that were sensitive to changes in odor concentration (ΔC_t_; Fig. 2). For these cell-odor pairs, responses to step stimuli differed from responses to static stimuli. These ΔC_t_ cells were selective for the direction of change, responding either to LH (Fig. 2a-b, S2d-f) or HL (Fig. 2c-d, S2a-c). For example, such a cell-odor pair may exhibit an identical response to static high and static low stimuli, but respond differently when these same concentrations are alternated in the HL stimuli (Fig. 2c-d). Such a response thus represents the concentration change rather than the concentration *per se*.

To categorize responses as ΔC_t_, CT, or CI, we tested whether the cumulative distribution of spike count after inhalation onset differed between stimuli (Kolmogorov-Smirnov test; Fig. 3a; Methods). A CI cell-odor pair responded identically to both concentrations, with or without a step (Fig. 3b1). A CT cell-odor pair responded differently to the two concentrations, and this difference is not affected by a concentration step (Fig. 3b2). If after a positive change in concentration, the cell responded differently from its response to static high concentration, this cell was categorized as +ΔC_t_. -ΔC_t_ cell-odor pairs gave a different response to the low concentration depending on the concentration in the preceding sniff. In summary, 51% (n=123) of cell-odor pairs responded to the odorants we presented. Of these responsive neurons, 41% were ΔC_t_, 20% were CT and 39% were CI (Fig. 3c). Cell-odor pairs that gave ΔC_t_ responses to one odor gave diverse responses to other odors presented in the same session (Fig. 3d). Thus, *ΔC_t_* sensitivity is not an odor-invariant property of these cells.

**Figure 3.**
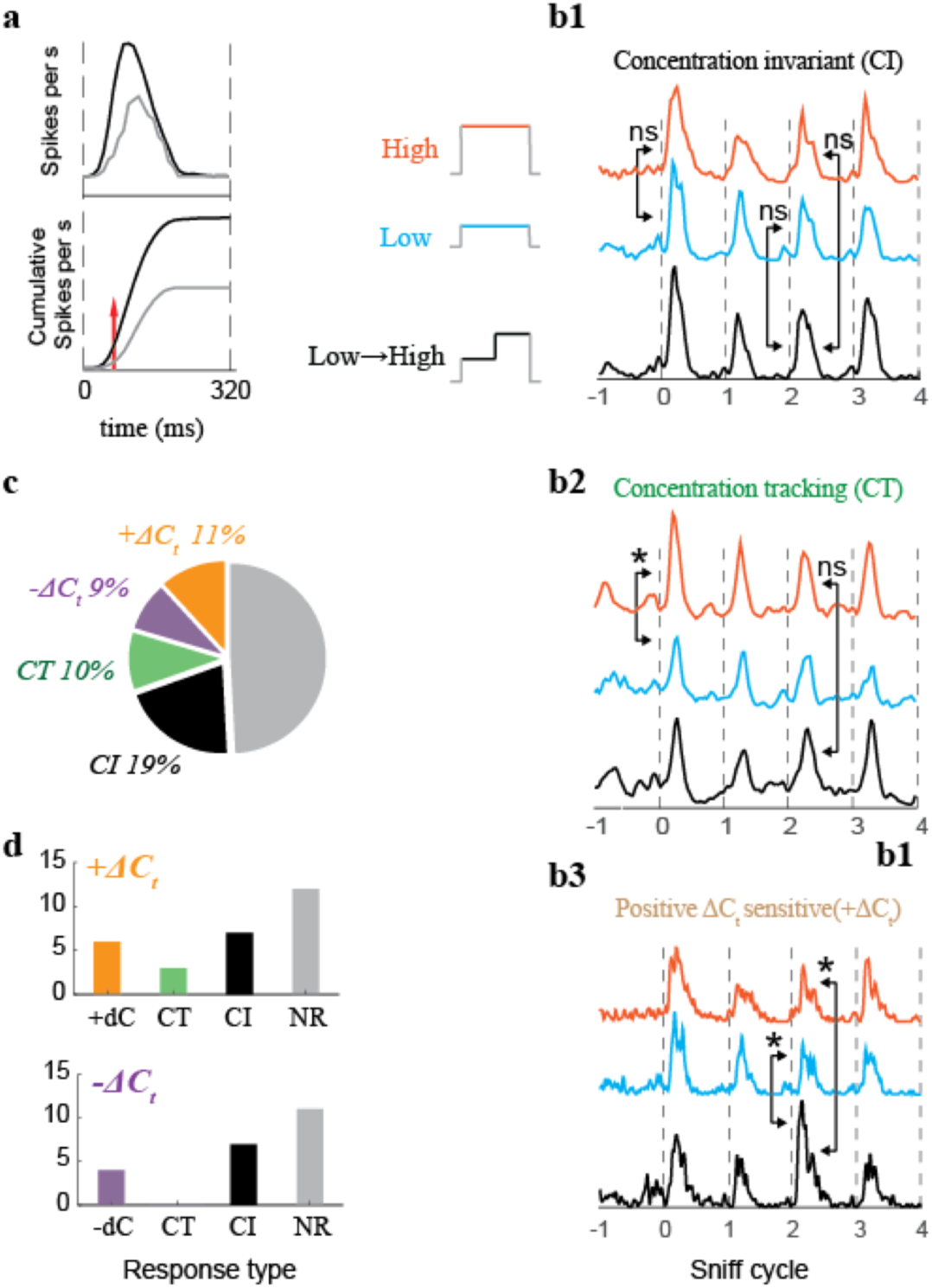
Categorization of response types. **a.** Criteria for determining whether a cell was responsive to a given odor. Top: Example of excitatory odor response PSTH. The black line is a PSTH of spiking during odorized sniffs. The grey line is a PSTH during unodorized sniffs. Bottom: cumulative spike counts of data from top plot. The red line indicates the first moment when cumulative distributions with and without stimulus become statistically different. **b1-3.** PSTHs from examples of each response type to high, low, and low->high stimuli are vertically separated. Arrows indicate which sniffs of the response are statistically compared. Non-significant differences are marked ns, and significant differences are marked with * (Kolmogorov-Smirnov test, p<0.01). **b1.** To be scored as concentration invariant (CI), the cell’s response on the first sniff of the static stimuli (high and low) must not significantly differ. In addition, the response on the third sniff of the concentration step stimulus must not differ from the third sniff of the two static stimuli. Example data are the same as Fig 1f. **b2.** Concentration tracking responses must differ on the first sniff of the static stimuli, but must not differ between the third sniff of step and static stimuli. Example data are the same as Fig. 1d. **b3.** ΔC_t_ sensitive responses must differ on the third sniff of the ΔC_t_ stimulus from the third sniff of both control stimuli. Only positive ΔC_t_ (+ΔC_t_) response is shown, but the rule is the same for negative ΔC_t_. Example data are the same as Fig 2b. **c.** Distribution of different response types: Concentration Invariant (CI; n=49), Concentration Tracking (CT; n=25), Positive ΔC_t_ (+ΔC_t_, n=28), and Negative ΔC_t_ (-ΔC_t_; n=21). **d.** Distribution of responses to a second odor for positive (top) and negative (bottom) ΔC_t_ cell-odor pairs.

### Contrast between concentrations depends on the stimulation history

In ΔC_t_ responses (Fig. 2), the response to a given concentration depends on the concentration presented in the previous sniff. This single sniff history dependence increases the difference between responses to different concentrations, thus enhancing the contrast. Responses of M/T cells may encode odor stimuli either by changes in spike count or by changes in temporal profile without changes in spike count ^18,19^. Our method of classifying responses is sensitive not only to changes in the total number of spikes within a sniff cycle but also to temporal redistribution of spikes within the cycle. To separately quantify which features of neuronal responses contribute to contrast enhancement, we compared the difference between responses to high and low concentrations when preceded by a step to the difference when preceded by the same concentration (Fig. 4a). We plotted full sniff spike count differences between the 3^rd^ sniffs of the two static stimuli (|High - Low|, Fig. 4b) against spike count differences between a dynamic step stimulus and the corresponding static stimulus (i.e., |Dynamic - Static|, Fig. 4b). In this visualization, the farther a cell-odor pair is from the diagonal, the stronger its contrast enhancement. Both +ΔC_t_ and -ΔC_t_ response populations showed contrast enhancement, with responses significantly shifted from the diagonal (t-test, P < 0.001, n=49), while the distributions for CT (t-test, P = 0.30, n=25) and CI (t-test, P = 0.18, n = 49) responses are symmetric about the diagonal.

**Figure 4.**
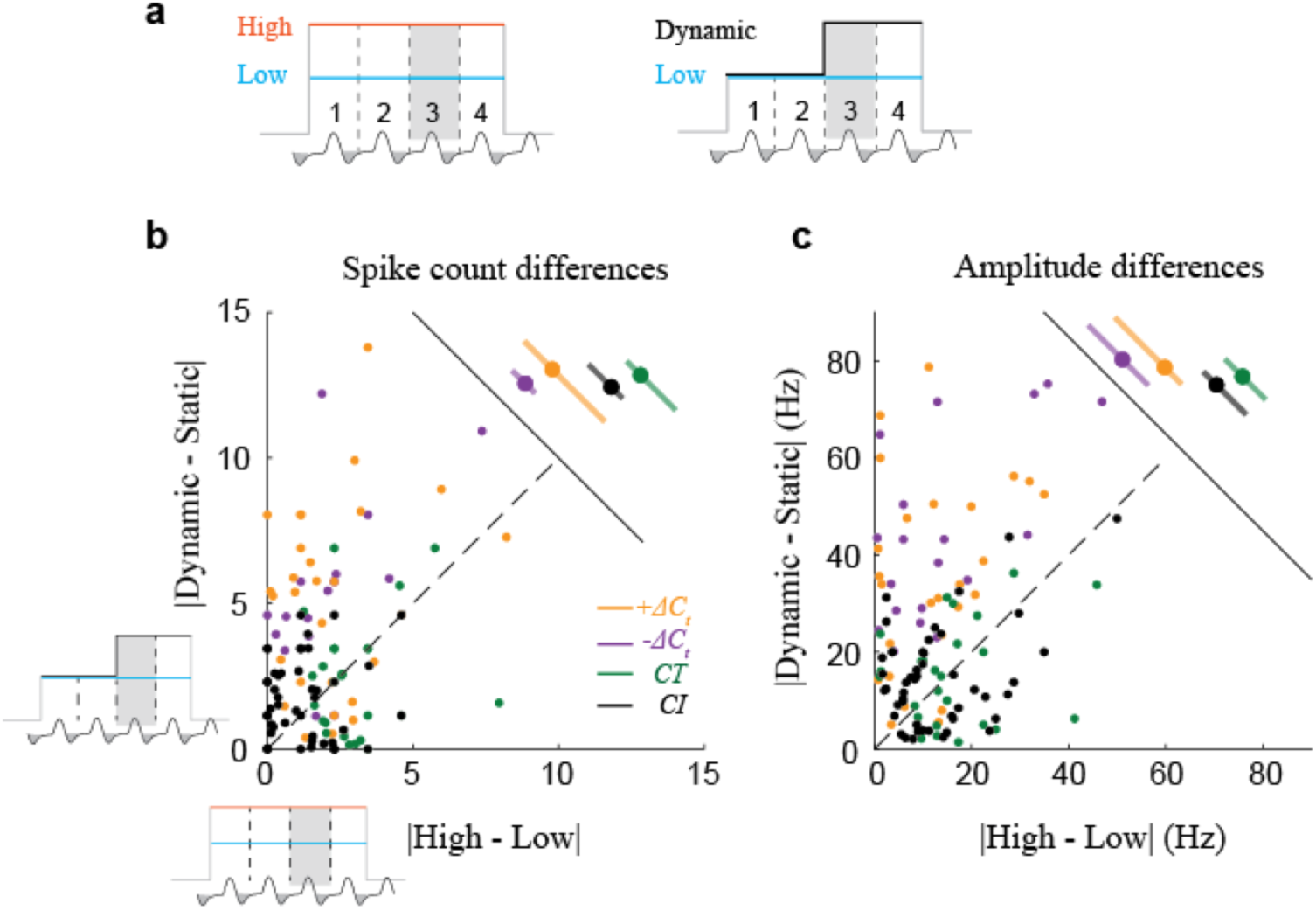
Contrast between concentrations depends on the stimulus history. **a.** Schematic of contrast comparison. To compare contrasts, for each cell-odor pair, we take the difference in response between the 3^rd^ sniffs of the static high (H) and static low (L) stimuli, and plot that against the difference between the 3^rd^ sniffs of the dynamic stimulus and the corresponding static stimulus (in this example L). Thus, only the concentration in the preceding sniff varies, and the concentrations being compared are constant. **b.** Scatter plot of full sniff spike count differences between two static stimuli against differences between dynamic and static stimuli, on 3^rd^ sniff cycle. CI, CT, +ΔC_t_ and -ΔC_t_ marked by black, green, orange, and blue color, respectively. Adjacent panel shows the means and STDs of the spike count differences. **c.** Same as **b** for differences in amplitude of the peak of the evoked PSTH. Adjacent panel shows the means and STDs of the peak amplitude differences.

To quantify how ΔC_t_ sensitivity enhances sub-sniff temporal differences between odor responses, we next performed the same comparison for differences in peak amplitude (Fig. 4c), a feature that reflects fast temporal patterning ^18,19^. Peak amplitude difference distributions for ΔC_t_ responses were significantly shifted from the diagonal (t-test, P < 0.01 for +ΔC_t_ and -ΔC_t_ responses), while for CT and CI responses the distributions were symmetric about the diagonal (t-test, P = 0.80 and 0.19, respectively). Thus, ΔC_t_ sensitivity also increased contrast at the faster sub-sniff timescale. Lastly, to determine the trial by trial reliability of contrast enhancement by ΔC_t_ responses, we used receiver operator characteristic (ROC) analysis (see Methods). In this analysis, ΔC_t_ responses discriminated better between dynamic and static stimuli than between two static stimuli (Fig. S3). These analyses demonstrate that ΔC_t_ sensitivity enhances the contrast between concentrations, potentially facilitating detection of concentration changes.

### ΔC_t_ sensitivity is step size dependent

We next tested how ΔC_t_ sensitivity depends on the size of the concentration step. In addition to the twofold steps used in the experiments above, we included a 1.5-fold and a 1.25-fold step, both LH and HL (Fig. 5a, d). To quantify the magnitude of ΔC_t_ sensitivity, we took the ratio of the response to the dynamic stimulus to that of the static stimulus, for full sniff spike count as well as peak amplitude of the PSTH. +ΔC_t_ responses (Fig. 5b) were largest for the 2-fold concentration increase, as expressed by the ratio of the response to the 3^rd^ sniff of the dynamic stimulus (LH_3_) to that of the corresponding static stimulus (H_3_), both for spike count and peak amplitude (Fig. 5c). Across the population of +ΔC_t_ responses, the two larger steps gave significant increases in spike count (t-test; 1.25-fold change: P=0.72; 1.5-fold change: P<0.01; 2-fold change: P<0.01), whereas only the largest step evoked a significant increase in peak amplitude: count (t-test; 1.25-fold change: P=0.5; 1.5-fold change: P=0.06; 2-fold change: P<0.001). For -ΔC_t_ responses (Fig. 5d-f), spike counts were significantly reduced for all step sizes tested (t-test; 1.25-fold change: P<0.01; 1.5-fold change: P<0.001; 2-fold change: P<0.01), while peak amplitudes were significantly reduced for the two larger steps (t-test; 1.25-fold change: P=0.019; 1.5-fold change: P<0.001; 2-fold change: P<0.001). Thus, the magnitude of contrast enhancement scales with the size of the concentration step.

**Figure 5.**
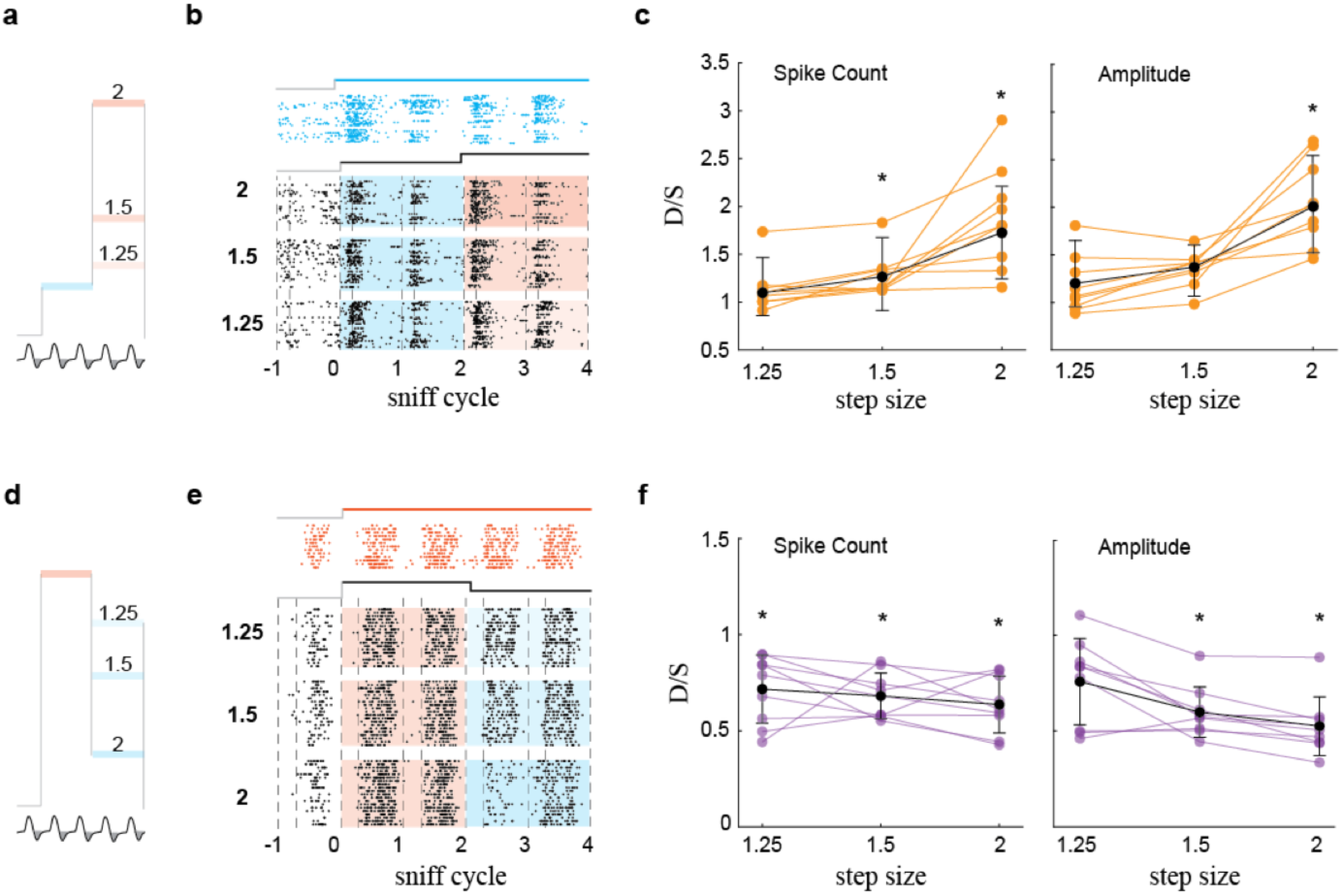
Contrast enhancement is proportional to the size of concentration change step. **a.** Stimulation with positive steps of different size. **b.** Raster plots of M/T cell’s activity during L static and three LH dynamic step stimuli. **c.** Normalized changes in spike count and amplitude of the response as function of step size. Orange lines are normalized changes for specific cell-odor pair, black line is the mean+/-std change across all responsive cell-odor pairs. Asterisks mark statistically significant deviation from 1 (t-test). **d-f.** Same for negative steps.

### Concentration decoding depends on temporal pattern, while ΔC_t_ decoding does not

M/T cell activity carries information about odor identity ^18-20^ and intensity ^21^ at sub-sniff timescales. To compare how information about concentrations and about changes in concentration might be decoded by downstream olfactory areas, we performed discriminant analysis (Methods). We first evaluated the accuracy with which responses to two odor concentrations can be discriminated by cell-odor pairs with a ΔC_t_ response (Fig. 6a). Classification of concentrations was performed on concatenated vectors of firing rates with multiple bin sizes: 5, 10, 20, 40, 80 and 160 ms. Concentration classification performance depended on bin size: smaller bin sizes yielded better discrimination (Wilcoxon signed-rank test; P < 0.01; Fig. 6c). Thus, information about odor concentration can be read out most accurately from fine timescale temporal patterns. Using the same classification procedure, we next evaluated whether decoding of concentration changes by the same ΔC_t_ cell-odor pairs similarly depends on temporal resolution (Fig. 6b). This analysis indicates that decoding of concentration changes is invariant across the full range of bin sizes (Wilcoxon signed-rank test, P = 0.14 Fig. 6c). These findings suggest that downstream neurons decode concentration and ΔC_t_ via different mechanisms.

**Figure 6.**
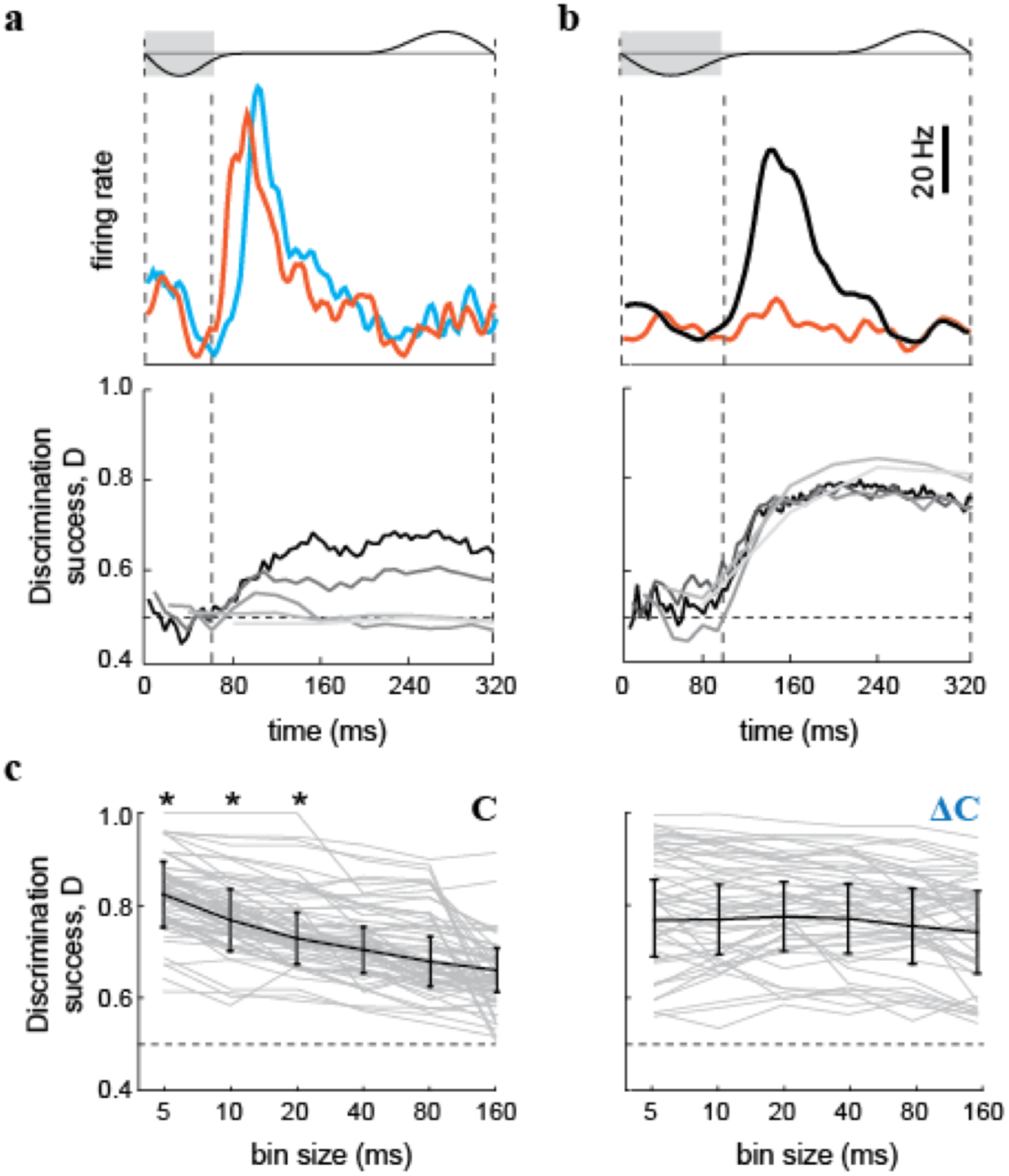
Discrimination among concentrations and changes in concentration by individual M/T cells. **a.** Top: PSTHs for a neuron’s responses to two static stimuli (red: high concentration, blue: low concentration). Bottom: Corresponding static stimuli discrimination success as a function of time. Vertical dashed lines indicate the end of the inhalation interval. Horizontal dashed lines indicate chance level performance. Different colored traces indicate discrimination success for different bin sizes. **b.** Top: PSTHs for a neuron’s responses to a high concentration static stimulus (red), and to a positive concentration step (black). Bottom: Corresponding static stimulus vs step stimulus discrimination success as a function of time. Different colored traces indicate discrimination success for different bin sizes. **c.** Discrimination performance of a linear classifier between two odor concentrations (left) and between changes in concentration (right) over the 320 ms window, as a function of bin size. Grey lines are performances of individual neurons. Black line is mean +/- std. Asterisks mark statistically significant deviations from discrimination success at 160 ms bin size.

## Discussion

Our work provides the first evidence that neurons in the mammalian olfactory bulb detect inter-sniff changes in odor concentration, thus enhancing temporal contrast. Such temporal contrast enhancement is widespread in other sensory modalities, consistent with the paramount importance of sensing stimulus dynamics.

Processing of odor dynamics works differently in invertebrate olfactory systems, because the olfactory organs of invertebrates continuously sample incoming odors. In these systems, olfactory neurons may represent gradients of odor concentration ^12,15,22^, as well as intermittent intensity fluctuations found in plumes ^23^. In contrast, terrestrial vertebrates such as mice sample odors intermittently. In order to compare the intensities of the previous and the current inhalation, the animal must preserve a representation of the previous concentration during the exhalation interval. This delayed comparison is likely implemented by intrabulbar circuits ^24^ or cortical feedback to the bulb ^25,26^, though we cannot exclude the possibility that it may be implemented in the OSNs.

Odor concentration gradients are critical for odor source localization. Vertebrates sense gradients by stereo (inter-naris) and serial (inter-sniff) comparisons ^7,27^. Because the nares are close together, stereo comparison should be most informative near an odor source, where odor gradients are steep. Shallower gradients, farther from a source, require the inter-sniff comparison, since the distance between sampling locations can be larger than the inter-naris distance ^7^. In a turbulent environment with noisy gradients^28^, comparison over more than two sniff cycles may be required. While stereo comparisons have been studied both behaviorally ^7,9,27^ and electrophysiologically ^27,29^, the serial component, which should dominate over a wider range of distances, has not been explored. Our study demonstrates a neural representation of ΔC_t_ detection. We propose that this representation contributes to odor source localization in natural olfactory scenes.

## Acknowledgments

We would like to acknowledge Yashar Ahmadian and Avinash Bhalla for discussions, Wolfgang Kelsch, Shawn Lockery, David McCormick, Kathy Nagel, Shy Shoham and Mike Wehr for comments on the manuscript. This research was supported by the Israel Science Foundation grant 816/14 (R.S.), Marie Curie Career Integration grant 334341 (R.S.) and by grant from the Whitehall Foundation (M.S.).

## Methods

### Animals

Data were collected in seven C57BL/6J mice. Subjects were 8–16 weeks old males at the beginning of recordings and were maintained on a 12-h light/dark cycle (lights on at 8:00 p.m.) in isolated cages in animal facility. All animal care and experimental procedures were in accordance with a protocol approved by the University of Haifa and University of Oregon Institutional Animal Care and Use Committees.

### Surgery

Mice were anesthetized using isofluorane gas anesthesia, and a head plate and a pressure cannula were implanted. For sniffing cannula implantation, we drilled a small hole in the nasal bone, into which the thin 7-8 mm-long stainless cannula (gauge 23 capillary tubing, Small Parts) was inserted, fastened with glue, and stabilized with dental cement ^30^. A small craniotomy was performed above one of the olfactory bulbs, contralateral to the side of sniffing cannula implantation. The reference electrode was implanted in cerebellum. At the end of the procedure, the craniotomy was covered with a biocompatible silicone elastomer sealant (Kwik-cast, WPI). The mice were given 3 days after a surgery for recovery.

### Electrophysiological recording

Before recording began, the mice were first adapted to head fixation. Mice typically remained quiescent after 1–2 sessions of head fixation, after which recording sessions started. We presented 2-3 odors in a single session in pseudo-random sequence with an average inter stimulus interval of 7 s and stimulus duration of 1–2 s. Each odor was presented in four temporal patterns: 1) static high – high concentration (∼1-2% of saturated vapor pressure) of odor for 4 sniff cycles; 2) static low – low concentration (50% of high concentration level) for 4 sniff cycles; 3) a step from high to low – for the first two sniff cycles, concentration level was equal to the level of static high, after which the concentration stepped to the low concentration; 4) and a step from low to high – two sniff cycles of low concentration followed by two sniffs of high concentration. We controlled odor concentration using a custom-built concentration change manifold (CCM, see next section). Odor onsets and concentration changes were triggered at the beginning of the exhalation phase, which occur at positive-going zero crossings of the pressure signal. Since odor cannot enter the nose during exhalation, triggering by exhalation onset allows enough time for the odor stimulus to reach a steady state of concentration by the time the animal begins to inhale. One session usually lasted for 60–90 min and consisted of 300–400 trials.

### Odor delivery

For stimulus delivery, we used a custom eight-odor air dilution olfactometer, based on a previous design ^31^. When no odor was being presented to the mouse, a steady stream of clean air (1,000 ml/min) was flowing to the odor port. During odorant presentation, N_2_ flowed through the selected odorant vial. We used multiple odorants obtained from Sigma-Aldrich. The odorants were stored in liquid phase (diluted either 1:5 or 1:10 in mineral oil) in dark vials. We used acetophenone, amyl acetate, geraniol, ethyl acetate, S - limonene, methyl butyrate, menthone, methyl salicylate, pentyl acetate and vanillin as odorants. The odorant concentration delivered to the animal was reduced additional tenfold by air dilution, and homogenized in a long Teflon tube before reaching the final valve. After sufficient mixing and equilibration time, the dual three-way Teflon valve (SH360T042, NResearch) directed the odor flow to the odor port, and diverted the clean airflow to the exhaust. All air flows and line impedances were equalized to minimize the pressure transients resulting from odor and final valve switching. Time course of odor concentration was checked by Photo-Ionization Detector (200B mini-PID, Aurora Scientific). The concentration reached a steady state ∼ 40 ms after final valve opening^32^. Further, to change odor concentration, we passed stable odorized airflow through a concentration change manifold (Fig. 1a). Odor concentration changes were achieved by activating a pair of matching solenoids (LHQA2411220H; The Lee Company) which performed air dilution. For each pair of solenoids, one valve was connected to a vacuum channel and the other to a clean airflow channel. Solenoid activation in the vacuum channel diverted part of the odorized air, while solenoid activation in the air channel contributed an equal amount of flow back into the system. To maintain constant total airflow (Fig. S1b), the impedance of each air channel was matched to the impedance of the corresponding vacuum channel using manual needle valves R_1_˙._3_ (NV3H-1012-3-S; Beswick Engineering). To ensure that the temporal profile of odor concentration stabilized before inhalation began, we predominantly used odorants with higher vapor pressure^33^. For these high vapor pressure odorants, the stimulus reaches 95% of final concentration in 20-40 ms (Fig. S1a).

### Electrophysiology and sniff signal recording

We recorded M/T cell activity using acute 16- or 128-channel matrix array of Si-probes (a2x2-tet-3mm-150-150-121-A16, M4x8-5mm-Buz-200/300um, NeuroNexus). Cells were recorded in both ventral and dorsal mitral cell layers. The data were acquired using a 128-channel data acquisition system (RHD2000, Intan Technologies) at 20 KHz sampling frequency. To monitor sniffing, the intranasal cannula was connected to a pressure sensor with polyethylene tubing (801000, A-M Systems). The pressure was measured using a pressure sensor (24PCEFJ6G, Honeywell). The amplified output signal from the pressure sensor was recorded in parallel with electrophysiological data on one of the analog input channels.

### Spike extraction and data analysis

All of the analysis discussed below was done in Matlab (MathWorks). Electrophysiological data were filtered between 300 Hz – 5 KHz and spike sorted. For spike sorting we used software package written by Alex Koulakov ^19^.

### Temporal alignment of responses

For analysis, sniffing traces were down-sampled to 1 kHz, and filtered in the range of 0.5–30 Hz. The inhalation onset and offset were detected by zero crossings of a parabola fit to the minima of the pressure signal following the onset of the inhalation. Inhalation onset/offset was defined as the first/second zero crossing of the parabola ^19^. We defined two intervals: the first is from inhalation onset to inhalation offset and the second is the rest of the sniffing cycle, from the inhalation offset to the next inhalation onset. While the duration of the first interval is concentration independent, the duration of the second interval depends on the concentration of presented odor (Fig. S4). To compare neuronal responses across trials and concentrations, we morphed the inhalation part of the sniff cycle and corresponding spike train to the average one ^19^. The second part of the sniff cycle and corresponding neural activity were artificially matched to the average over trials: longer cycles were truncated and shorter were zero padded.

### Odor responses

To establish whether a cell is responsive to an odor, we compared the cumulative distribution of the neuronal spikes without odors to the cumulative distribution of the neuronal activity during first odorized sniff cycle, using the Kolmogorov-Smirnov test. Neuronal activity without odor was sampled from 3 sniffs preceding odor delivery across all trials. Neuronal activity for a given odor was sampled from the first sniff after stimulus onset. Cells were considered responsive if the distribution of spiking activity during the first odorized cycle statistically differed from the distribution of baseline responses in at least one 10 ms bin relative to inhalation onset (p < 0.005; Benjamini-Hochberg multiple comparison correction) or if their average spike rate over the sniff cycle differed significantly from baseline (p < 0.05).

For visualization purpose only, we estimated standard deviation of the peak amplitude of the responses in Figures 1c-f and 2. Single response amplitude was constructed from 70% of trials which were randomly selected. This procedure was repeated 200 times for different single trial population. Standard deviation was calculated using distribution of response amplitudes.

To categorize responses as ΔC_t_, CT, or CI, we tested whether the cumulative distribution of spike count after inhalation onset was significantly different between different stimuli on the 1^st^ and 3^rd^ sniff cycles. This method is sensitive not only to changes in total amount of spikes per sniff but also to temporal redistribution of spikes within the cycle. Two conditions had to be fulfilled to score a response as CI: 1) no significant differences between responses to H and L stimuli on the 1^st^ sniff cycle; 2) no significant differences between responses to HL or LH and corresponding static stimulus on the 3^rd^ sniff cycle (Fig. 3b1). If the 2^nd^ condition is satisfied but the 1^st^ is not, then the response was scored as CT (Fig. 3b2). A cell-odor pair is deemed as ΔC_t_ if its response on the 3^rd^ sniff cycle differs significantly from responses to both H and L stimuli on the same sniff cycle (Fig. 3b3). If after a positive change in concentration, the cell responds differently from its response to static high concentration, this cell is scored as +ΔC_t_. -ΔC_t_ cell-odor pairs gave a different response to the low concentration depending on the concentration in the preceding sniff.

### ROC analysis

ROC analysis provides a measure of how well a given cell-odor pair can discriminate between two stimuli. To measure the discriminability between the static odor stimuli, high and low, we first compute the area under the ROC curve (auROC) for the distributions of spike counts over the first sniff of each stimulus. We next compute the auROC between the sniffs before and after a concentration step in a LH and HL stimulus (Fig. S3a). We then plot the static stimulus discriminability against the ΔC_t_ discriminability. This plot shows whether a given cell-odor pair shows contrast enhancement between concentrations during step stimuli. In our ROC based discriminability index, a value of 1 indicates no overlap between the two distributions, and perfect discriminability in ΔC_t_, while a value of 0.5 indicates complete overlap between the two distributions and inability to discriminate ΔC_t_.

The three example cell-odor pairs are shown in such a plot (Fig. S3). Concentration invariant responses do not discriminate between high and low concentration, and have values of near 0.5 for both static and step stimuli. Concentration-tracking responses discriminate between step stimuli and corresponding control equally as well as they discriminate between the two static control stimuli. Thus, they fall along the diagonal of this plot. Finally, ΔC_t_ responses discriminate better between sniffs of step stimuli than for sniffs of static stimuli, so they fall above the diagonal.

### Odor concentrations classification analysis

To estimate how well single neurons (n=49) can discriminate between two odor concentrations on a trial by trial basis, we constructed a Mahalanobis distance linear classifier. For concentration discrimination, we calculated discriminability between responses to static high and static low on the 3^rd^ sniff cycle, L_3_ and H_3_. For every cell and for every pair of concentrations we counted spikes using multiple time bins (5, 10, 20, 40, 80 and 160 ms). Single trials were randomly selected and compared to a set of templates constructed from 70% of trials for each of the two concentrations. We used the *mahal* function in Matlab to estimate Mahalanobis distance from each single trial vector to two groups of multiple trial templates representing two concentrations. This procedure was repeated 300 times for different single trial population vectors and was repeated for each bin size.

A similar analysis was performed on the same cell-odor pairs to estimate discriminability in ΔC_t_. For ΔC_t_ discrimination we calculated discriminability between LH_3_ and L_3_ sniffs for +ΔC_t_ responses and HL_3_ and H_3_ sniffs for -ΔC_t_ responses.

## Data availability

Request for materials should be addressed to R.S.

## References

1. Wertheimer, M. Experimentelle Studien über das Sehen von Bewegung. (1912).

2. Bregman, A. S. Auditory Scene Analysis. (MIT Press, 1994).

3. Werner, G. & Mountcastle, V. B. Neural activity in mechanoreceptive cutaneous afferents: stimulus-response relations, Weber functions, and information transmission. Journal of Neurophysiology 28, 359–397 (1965).

4. Ratliff, F., Hartline, H. K. & Miller, W. H. Spatial and temporal aspects of retinal inhibitory interaction. J Opt Soc Am 53, 110–120 (1963).

5. Langner, G. Periodicity coding in the auditory system. Hear. Res. 60, 115–142 (1992).

6. Mountcastle, V. B., Talbot, W. H., Darian-Smith, I. & Kornhuber, H. H. Neural basis of the sense of flutter-vibration. Science 155, 597–600 (1967).

7. Catania, K. C. Stereo and serial sniffing guide navigation to an odour source in a mammal. Nature Communications 4, 1441 (2013).

8. Khan, A. G., Sarangi, M. & Bhalla, U. S. Rats track odour trails accurately using a multi-layered strategy with near-optimal sampling. Nature Communications 3, 703 (2012).

9. Porter, J. et al. Mechanisms of scent-tracking in humans. Nat. Neurosci. 10, 27–29 (2007).

10. Fujiwara, T. et al. Odorant concentration differentiator for intermittent olfactory signals. Journal of Neuroscience 34, 16581–16593 (2014).

11. Larsch, J. et al. A Circuit for Gradient Climbing in C. elegans Chemotaxis. CellReports 12, 1748–1760 (2015).

12. Kim, A. J., Lazar, A. A. & Slutskiy, Y. B. Projection neurons in Drosophila antennal lobes signal the acceleration of odor concentrations. eLife 4, (2015).

13. Vickers, N. J. Mechanisms of animal navigation in odor plumes. Biol. Bull. 198, 203–212 (2000).

14. Huston, S. J., Stopfer, M., Cassenaer, S., Aldworth, Z. N. & Laurent, G. Neural Encoding of Odors during Active Sampling and in Turbulent Plumes. Neuron 88, 403–418 (2015).

15. Dynamical feature extraction at the sensory periphery guides chemotaxis. eLife 4, (2015).

16. Cleland, T. A. et al. Sequential mechanisms underlying concentration invariance in biological olfaction. Front Neuroeng 4, 1–12 (2011).

17. Wilson, R. I. & Mainen, Z. F. Early events in olfactory processing. Annu. Rev. Neurosci. 29, 163–201 (2006).

18. Cury, K. M. & Uchida, N. Robust Odor Coding via Inhalation-Coupled Transient Activity in the Mammalian Olfactory Bulb. Neuron 68, 570–585 (2010).

19. Shusterman, R., Smear, M. C., Koulakov, A. A. & Rinberg, D. Precise olfactory responses tile the sniff cycle. Nat. Neurosci. 14, 1039–1044 (2011).

20. Bathellier, B., Buhl, D. L., Accolla, R. & Carleton, A. Dynamic ensemble odor coding in the mammalian olfactory bulb: sensory information at different timescales. Neuron 57, 586–598 (2008).

21. Sirotin, Y. B., Shusterman, R. & Rinberg, D. Neural Coding of Perceived Odor Intensity. eNeuro 2(6), 10.1523/ENEURO.0083 (2015).

22. Nagel, K. I. & Wilson, R. I. Biophysical mechanisms underlying olfactory receptor neuron dynamics. Nat. Neurosci. 14, 208–216 (2011).

23. Vickers, N. J., Christensen, T. A., Baker, T. C. & Hildebrand, J. G. Odour-plume dynamics influence the brain's olfactory code. Nature 410, 466–470 (2001).

24. Sturm, T. et al. Mouse urinary peptides provide a molecular basis for genotype discrimination by nasal sensory neurons. Nature Communications 4, 1616 (2013).

25. Luskin, M. B. & Price, J. L. The topographic organization of associational fibers of the olfactory system in the rat, including centrifugal fibers to the olfactory bulb. J. Comp. Neurol. 216, 264–291 (1983).

26. Li, Z. A model of olfactory adaptation and sensitivity enhancement in the olfactory bulb. Biol Cybern 62, 349–361 (1990).

27. Rajan, R., Clement, J. P. & Bhalla, U. S. Rats smell in stereo. Science 311, 666–670 (2006).

28. Murlis, J., Elkinton, J. S. & Carde, R. T. Odor plumes and how insects use them. Annual review of entomology 37, 505–532 (1992).

29. Kikuta, S. et al. From the Cover: Neurons in the anterior olfactory nucleus pars externa detect right or left localization of odor sources. Proc Natl Acad Sci USA 107, 12363–12368 (2010).

30. Verhagen, J. V., Wesson, D. W., Netoff, T. I., White, J. A. & Wachowiak, M. Sniffing controls an adaptive filter of sensory input to the olfactory bulb. Nat. Neurosci. 10, 631–639 (2007).

31. Bodyak, N. & Slotnick, B. Performance of mice in an automated olfactometer: odor detection, discrimination and odor memory. Chemical Senses 24, 637–645 (1999).

32. Resulaj, A. & Rinberg, D. Novel Behavioral Paradigm Reveals Lower Temporal Limits on Mouse Olfactory Decisions. Journal of Neuroscience 35, 11667–11673 (2015).

33. Martelli, C., Carlson, J. R. & Emonet, T. Intensity invariant dynamics and odor-specific latencies in olfactory receptor neuron response. Journal of Neuroscience 33, 6285–6297 (2013).

